# Reducing variance in a mouse defect healing model: Real-time Finite Element Analysis allows homogenization of tissue scale strains

**DOI:** 10.1101/2020.09.02.274878

**Authors:** Graeme R. Paul, Esther Wehrle, Duncan C. Tourolle, Gisela A. Kuhn, Ralph Müller

## Abstract

Mechanical loading allows both investigation into the mechano-regulation of fracture healing as well as interventions to improve fracture-healing outcomes such as delayed healing or non-unions. However, loading is seldom individualised or even targeted to an effective mechanical stimulus level within the bone tissue. In this study, we use micro-finite element analysis to demonstrate the result of using a constant loading assumption for all mouse femurs in a given group. We then contrast this with the application of an adaptive loading approach, denoted real time Finite Element (rtFE) adaptation, in which micro-computed tomography images provide the basis for micro-FE based simulations and the resulting strains are manipulated and targeted to a reference distribution. Using this approach, we demonstrate that individualised femoral loading leads to a better-specified strain distribution and lower variance in tissue mechanical stimulus across all mice, both longitudinally and cross-sectionally, while making sure that no overloading is occurring leading to refracture of the femur bones.

## Introduction

Bone requires mechanical stimulation for fractures to heal. Improved understanding of the mechanical stimulation – fracture-healing relationship will provide substantial benefit in both basic scientific investigation of cell fate and behaviour as well as clinical application. However, the exact mechanical stimulation required to initiate bone formation is still up to debate, with *in vivo, in vitro* and *in silico* models showing differing levels of activation strains at different anatomical scales^1-4^. For example, at tissue level, strains of up to 3’000 microstrain occur due to strenuous physiological loading in humans ^5^, while at cell level, *in silico* models have shown that micro-architectural variations lead to strain amplifications and strain peaks exceeding 10’000 microstrain in and around osteocytes^6,7^. These levels of strain approach the activation levels seen in single cell *in vitro* investigations on the response of osteocytes to mechanical loading^8^. However, even though mechanical stimulation at each scale differs greatly, our ability to control the mechanical stimuli is most easily performed at organ scale, manipulating the bone via some sort of actuation resulting in a specific strain distribution at each scale. Controlling this “mechanical dose” in the mechanical environment is essential for experimental methods to either optimise or map mechanical stimulation to understand the formation processes underlying bone healing and develop improved interventions. In turn, with the growth of personalised medicine approaches, treatments adjusted towards a patient’s individual anatomy would require mechanical interventions with a specific “mechanical dose” for each patient.

The mechanical environment is greatly dependant on the geometry of bone^9^. Strains propagate differently through each individual bone, providing a variety of mechanical stimuli throughout the tissue^10^. This geometric variance is seen from traumatic fractures in humans^11^ to well controlled osteotomies in sheep^12^ or murine models^13^. While inbred mouse or rat strains should provide the lowest variance out of all appropriate model animals for fracture healing, often the variance is not as low as expected. For example, even within studies, defect size variance can be large, often displaying a standard deviation of over 10% of the nominal size in osteotomy based experiments^12-14^. In addition, even though it is known that the resulting strain from the initial size of the fracture gap^12,15-18^ is influential in the outcome of the healing process, this geometric information is regularly left either unreported, with no size or geometric description of the defect presented^19-21^, or underreported with only the nominal size being presented^22-24^. While such basic geometric conditions are critical for outcome, additional affecters such as activity of the animal^25^, disruption of the periosteum^26,27^ and inter-fragmentary movement^28^ provide further biological and mechanical cues. These cues influence the longitudinal progression of bone formation and resorption and hence lead to geometric changes, which in turn further influence the mechanical environment within the tissue and the resultant healing outcome divergence across a group. Additionally, effects attributed to interventions such as pharmacological agents could be obscured due to inconsistent mechanical environments within and between groups. As loading models are often used to study the effects of mechanical stimulation on physiological processes in bone such as remodelling and healing, controlling the local and global stimuli acting on the tissue would improve the validity of such investigations^16^.

Non-individualised loading has been applied during all phases of the fracture healing process. Vibration loading has often been attempted during the inflammation phase with mixed results ^20,23,24,29^, while results that are more consistent have been seen with direct mechanical stimulation during the end stages of the reparative phase and the remodelling phase ^30,31^. However, while these studies aim at providing some degree of mechanical stimulation to the bone structure, limitations lie in the lack of either targeted mechanical stimulation (i.e. attempting to achieve a certain mechanical strain within the tissue), or application of load regardless of the individual structural and geometric state of the bone ^32-34^. Often attempts at targeting strains and the resultant mechanical loads are derived from past studies using bone-surface strain gauge measurements, amalgamating loading-strain relationships throughout the animal group ^35,36^. However, strain gauged based mechanical loading values show substantial in-group variance, with values differing by up to 30% in such studies ^35,36^.

In this study, we analyse the inter-individual and temporal variance of the mechanical environment using image-based finite element analysis ^16^ during the bridging and post-bridging phase under a 10 N load and, due to individual differences seen within groups, propose and apply a novel methodology of adapting loading conditions to the individual bone geometry within an *in vivo* mouse femur osteotomy model. We term this method real time Finite Element (rtFE) adaptive loading. Via the incorporation of finite element simulations into the experimental pipeline in real time, we are able to homogenise tissue level strains between each mouse, adapting the experimental load based on these simulations in one imaging and loading session. Additionally, we are able to assess whether a bone has a risk of refracture when under loading, allowing loading on the healing mouse femur to start as soon as it is safe to do so. This allows complete intervention control throughout the repair and remodelling phases of fracture healing.

## Results

In this study, we compared the effect of traditional constant loading (Fig. 1a) with loading adjusted by rtFE adaptive loading on the mechanical environment in a healing bone defect (Fig. 1b, c). We used longitudinal, time-lapsed *in vivo* micro-CT images (5 per animal) taken from 30 mice during healing of a femur osteotomy fixed by an external fixator. Ten of the animals received loading of the defect, the rest were sham loaded at 0 N. Based on the CT images, we calculated the local strain distribution within the callus by micro-FE analysis for a simulated 10 N loading. In a second step, we adapted the applied loading such that the distribution of strains is minimised with reference to a target distribution, derived from a mouse displaying a good healing progression. The whole process was optimized to allow incorporation into a single anaesthesia session, hence the name real time FE. We successfully ran the rtFE adaptive loading process with an increase of less than 5 min, in addition to image reconstruction times, between the end of scanning and the start of loading (Fig. 1c). This is important as it prevents the need to re-anaesthetise the animal, which induces stress and could influence study results.

**Figure 1:**
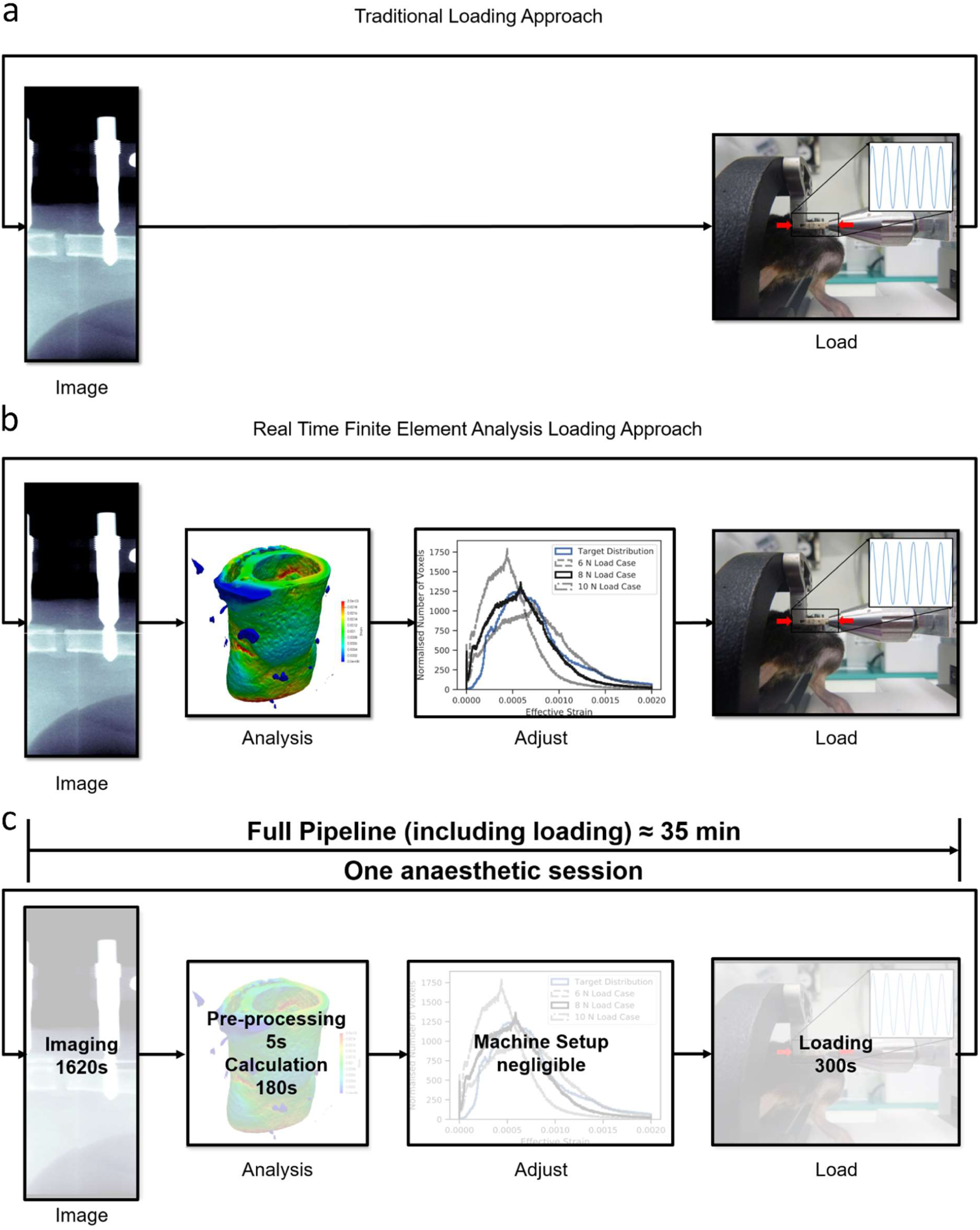
a) A traditional loading experiment images the animal and uses a loading protocol decided on before the experiment has begun. b) The rtFE approach incorporates the simulation and adjustment of the loading parameters to ensure a targeted mechanical stimulation at each time point for the animal. c) When incorporated within the experimental pipeline, appropriate implementation of the rtFE will allow the imaging to loading process to be incorporated in a single anaesthetic session.

### Longitudinal observation of the bone healing process

Bridging of the fracture defect occurred between week 2 and week 3 post surgery. As seen in Fig 2a and 2d, between week 2 and week 3, a considerable amount of bone was formed. The callus structure had many small strut- and truss-like features that transfer the structural load. These can be seen at week 3 (Fig. 2a and Fig. 2b). These small structural elements concentrated mechanical strains into small regions, increasing strain variance, such as seen at week 2, 3 and week 4 in Fig 2b, c, e and f, where strain standard deviation exceeds future values. Once these small structural elements were absorbed into the new cortical wall (as can be seen in Fig 2a and b, week 3 to 5), the strain distributions displayed lower standard deviation, as the thicker structure dissipates the load more evenly.

**Figure 2:**
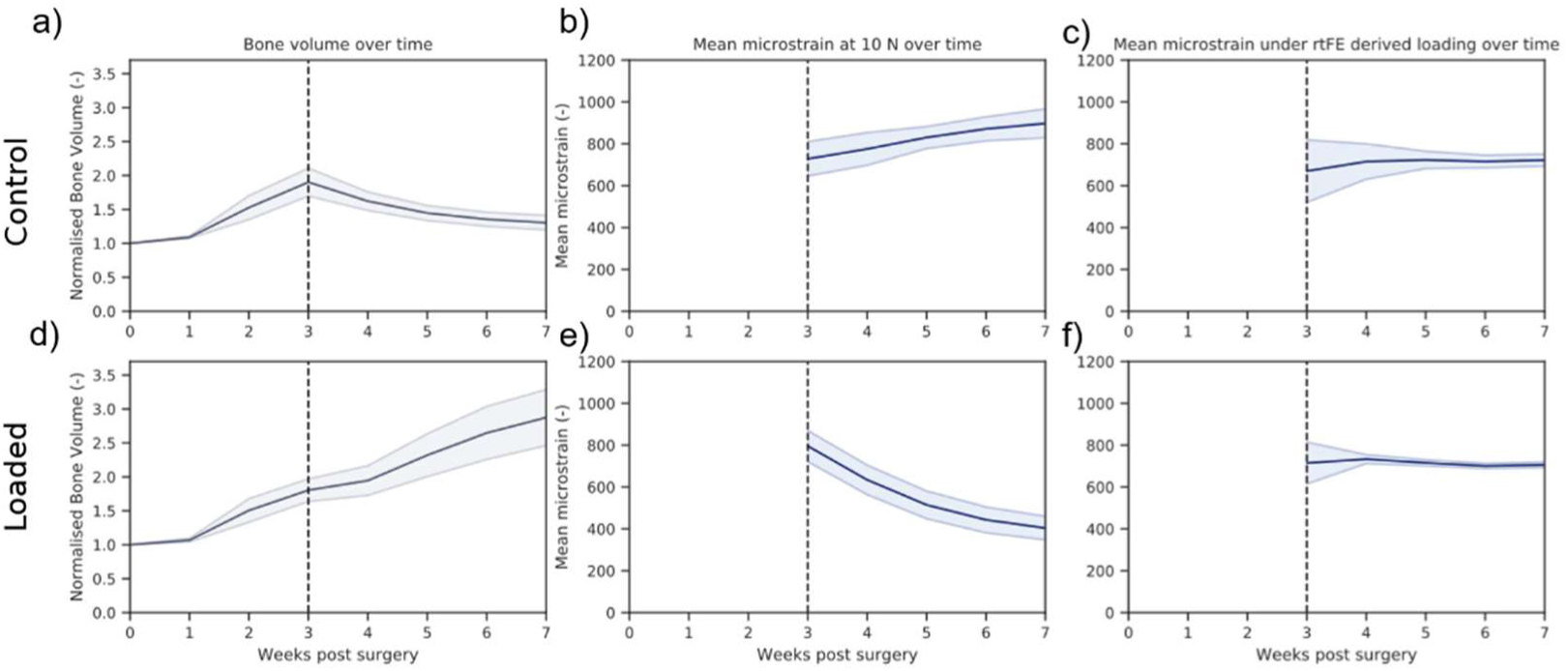
As bone volume decreases (a), or increases (d), the mechanical strains increase (b) or decrease (e) respectively throughout the duration of the loading period. This causes a changing mechanical environment at each point of time. (c) and (f) display the application of the rtFE approach to ensure this change in mechanical environment does not occur. Additionally comparisons between (b) and (c), and (e) and (f), show the reduction of variance caused by the rtFE specifying appropriate loading parameters on an individualised basis. Bone volume is normalised to the amount of bone volume at week 0 post surgery.

### Constant loading and the mechanical environment

At week 3 for the control group, with a load of 10 N, the median strain under constant loading was 683±81 microstrain. With the observed decreasing bone volume (Fig 2a), the median strain and standard deviation increased throughout the study duration, 740±60 microstrain at week 4, 818±61 at week 5, 873±71 at week 6 and 905±86 at week 7 (Fig 2b).

Contrastingly, in the loaded group, with an increasing bone volume (Fig 2d), the median strain decreased throughout the remaining reparative and remodelling phase (727±74 at week 3, 582±77 at week 4, 471±68 at week 5, 413±61 at week 6 and 383±56 at week 7) (Fig 2e). For both loading and control group, the standard deviation of strains in the mechanical environment remained high, roughly or greater than 10% of the median strain for all simulated time points.

### Real time Finite Element adaptive loading

Adaptation of loading parameters to ensure minimisation of strain variance was then performed, in a means that can be incorporated into an experiment; a process we termed real time Finite Element (rtFE) adaptive loading. In this process, finite element analysis is performed after the animal is imaged and the results are used to change the experimental loading parameters, ensuring similar strain distributions within or across groups. The rtFE adaptation required two stages: a strain distribution matching stage, to minimise variance between longitudinal and cross sectional mechanical environments, and a fracture-risk identification stage; where given the determined loading parameters, the risk of fracture was identified and the load downscaled if required.

A target distribution was developed from a mouse with good healing progression and scaled to be of a median strain of 700 microstrain. This acted as an idealised strain distribution for the application of rtFE. We then minimised the Kolmogorov-Smirnov test statistic (Fig 3a) between the mechanical environment of each mouse from week 3 to week 7 and this target distribution using a Nedler-Mead optimiser. This ensured minimal differences between the cumulative distribution functions of each distribution. Alternatively, the strain distribution can be plotted with a series of incremental possible loading options governed by whatever mechanical actuator is in use. The researcher can then select the most appropriate distribution (Fig 3b), where each possible scenario is plotted with regard to the target distribution.

**Figure 3:**
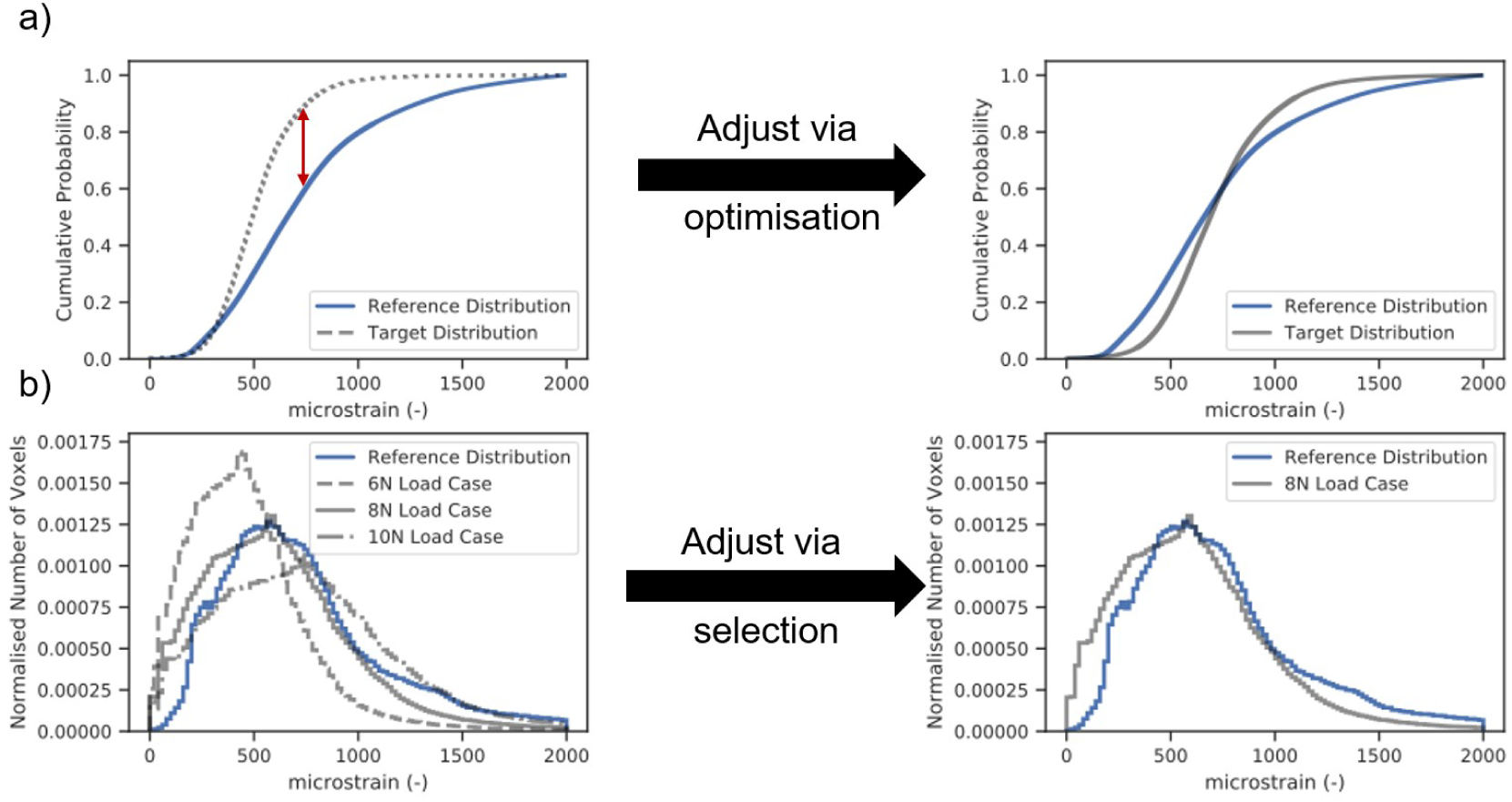
Longitudinal and cross-sectional strain progression under constant loading and rtFE adaptation simulations results show the progression of strain within the bone tissue. (a) demonstrates how poorly healed bones have less consistent strain distributions throughout the tissue, with regions of dangerously high strain and region of lower strains. Well-healed bones however show a decrease in the strains throughout the tissue, as is the case in L06. (b) rtFE adapts the loading to ensure no peak strain voxels (C10) and maintains consistent strain fields (L06). For mice with smaller changes in bone geometry (C09 and C14), the adaptation is less obvious, but still present.

However, scaling load and matching strain distribution can lead to certain fragile structures within the callus developing dangerously high strain, potentially leading to refracture. After the strain distribution matching is complete, a fracture risk analysis is performed. The aim of this analysis is to determine if the new loading parameters could lead to failure of the healing bone, and hence such an occurrence can be mitigated. For the selected loading case, the number of voxels of 10’000 microstrain or greater are counted. If this exceeds fifty voxels, the load is downscaled by 2 N and the number of voxels greater than 10’000 microstrain is counted again. This process is repeated until less than 50 voxels exceeding 10’000 microstrain remain (as depicted in Fig 4a-f) and the final loading parameter is selected.

**Figure 4:**
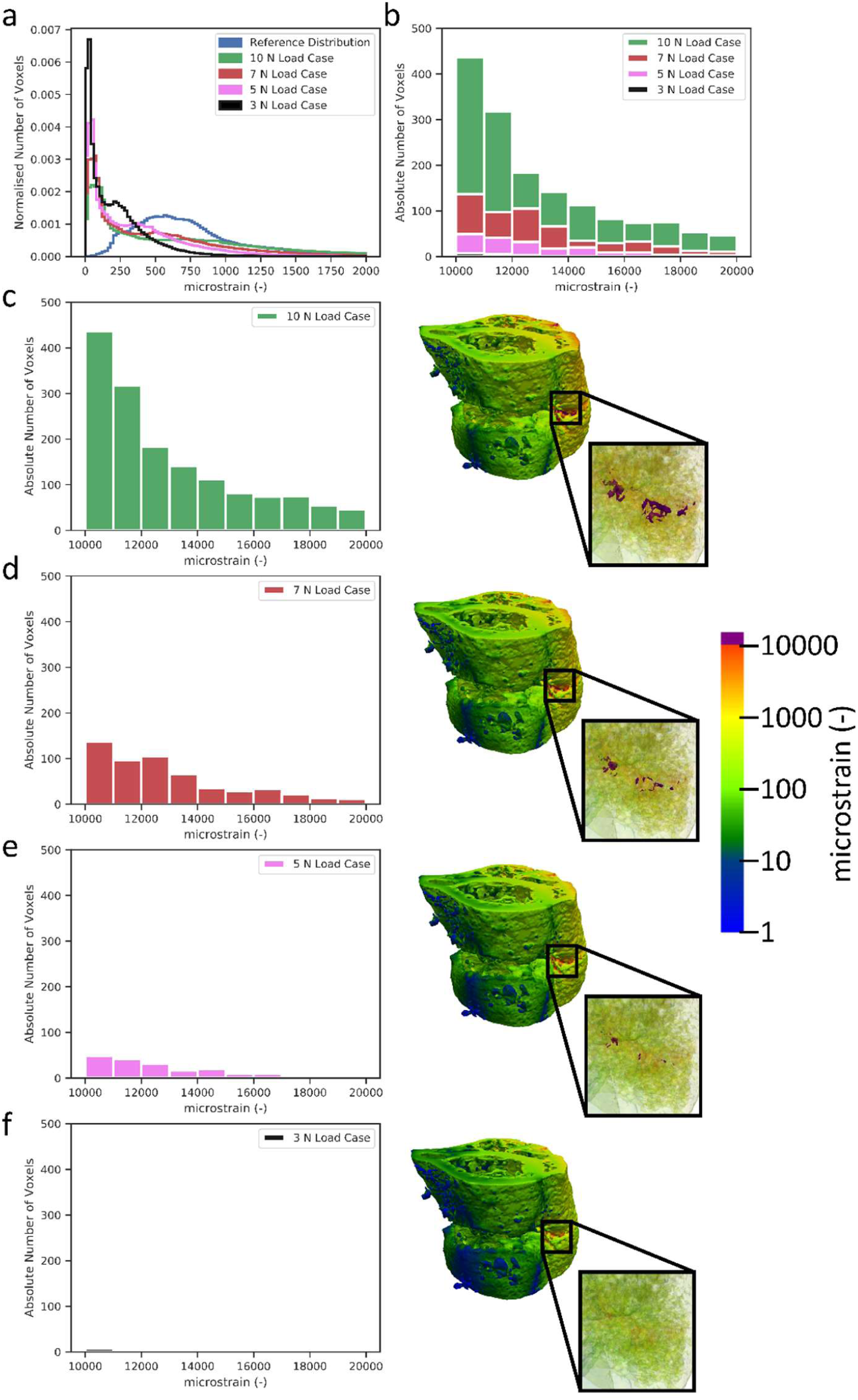
The fracture risk profiling (a) identifies the number of voxels over 1% strain (b) and down scales the load (c – f) until there are effectively no voxels in the identified strain risk region.

**Figure 5:**
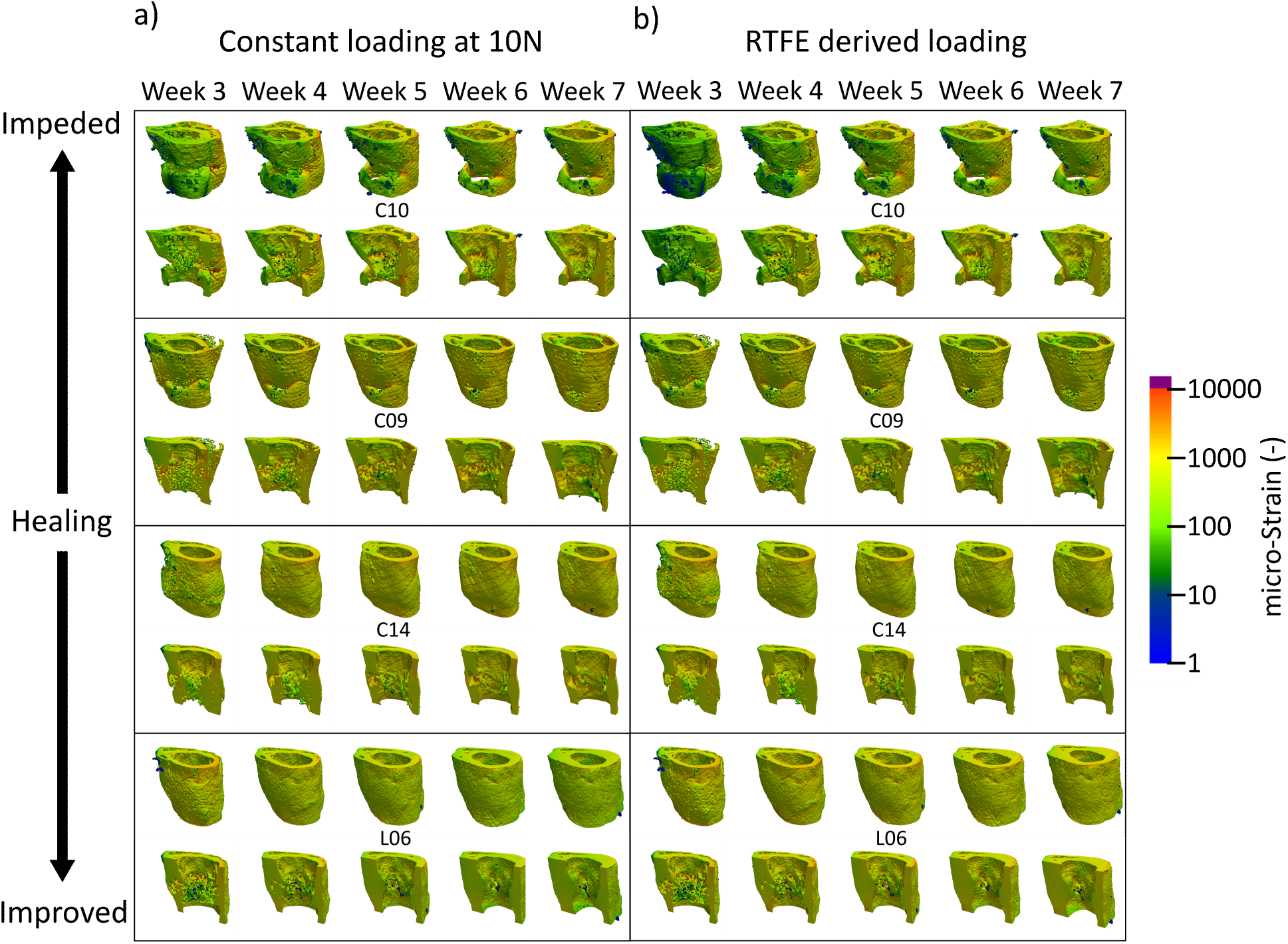
Two approaches can be implemented for the rtFE approach. Optimising via a Kolmogorov-Smirnov test (a) allows an automated process, saving time and giving an exact value, while plotting and selecting (b) from a list of loading options provides the researcher with more understanding of the strain distributions for each loading parameter.

### Adaptive loading and the mechanical environment

With application of rtFE adaptive loading (Fig 6), a set of loading parameters was determined. Control mice required a deceasing load throughout the observation period, while loaded mice required an increasing load. When looking at the mechanical environment, the median strain for the control group remained within narrow bounds (713±33 at week 3, 792±35 microstrain at week 4, 714±38 at week 5, 726±35 at week 6 and 736±30 at week 7) throughout the remaining reparative and remodelling phase, respectively. A similar pattern was seen in the loaded group with median strains remaining similar (721±32 at week 3, 712±20 microstrain at week 4, 700±15 at week 5, 681±14 at week 6 and 698±14 at week 7) from week 3 to week 7. As with the control group, upon rtFE application, standard deviation decreased substantially from week 3 to week 4, and further decreased during the remodelling period, remaining 35% and 25% of the constant load case for the control and loaded groups at week 7 respectively.

**Figure 6:**
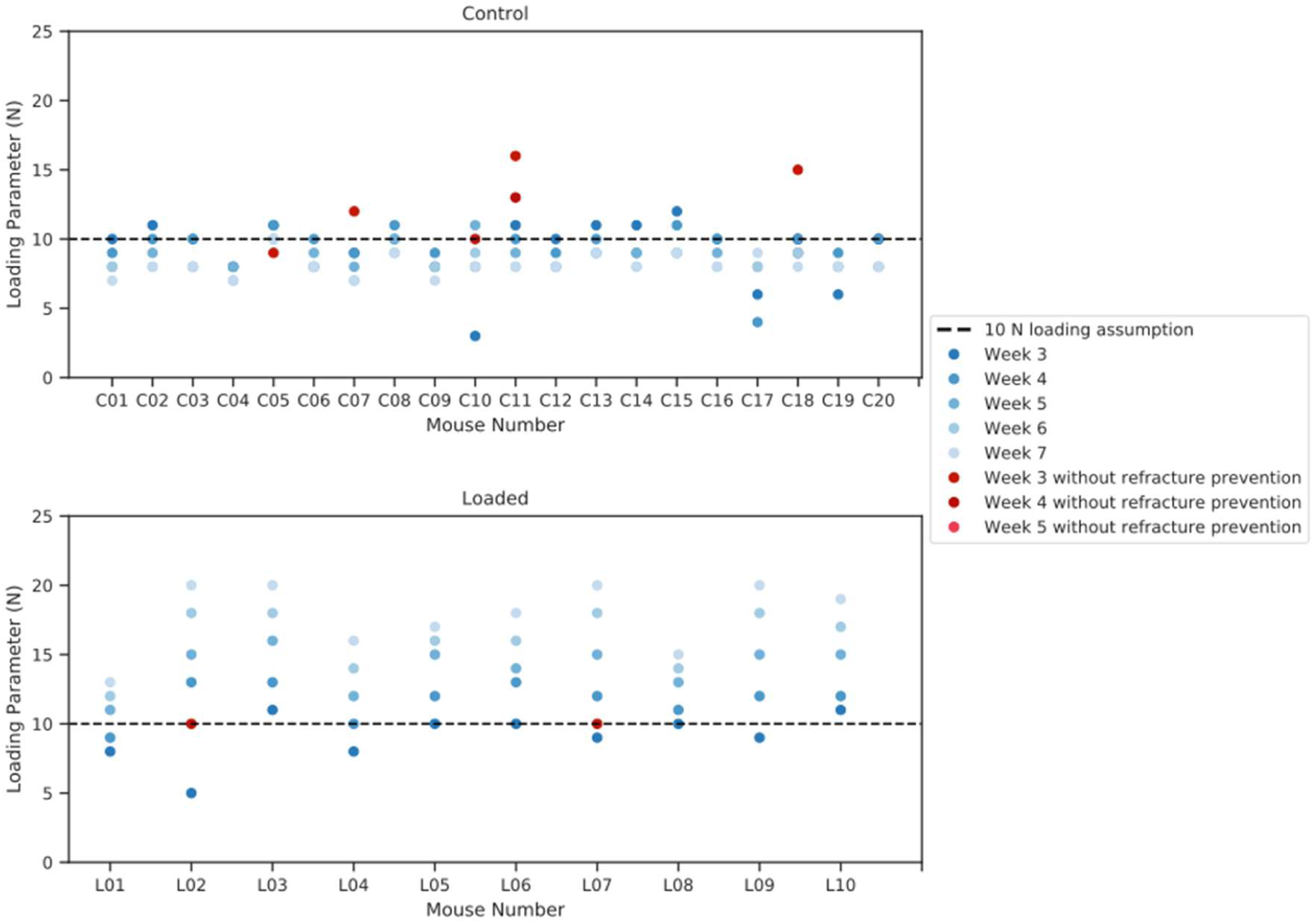
Loading parameters relative to the 10 N assumption, determined by rtFE. (a) a decreasing load is needed to maintain a consistent mechanical environment in a control group. (b) an increasing load is needed to maintain a consistent mechanical environment in a loaded group.

Upon application of the fracture prevention step, the control group observed a decrease in median strain and an increase in standard deviation (give mean increase/decrease in %; 691±158 at week 3, 726±110 microstrain at week 4, 714±47 at week 5, 726±44 at week 6 and 736±30 at week 7) (Fig 2c) in comparison with the mechanical environment prior to this step. Likewise, the loaded group also displayed an increase in standard deviation for earlier time points yet a quicker reduction than in the unloaded group (721±122 at week 3, 712±20 microstrain at week 4, 700±15 at week 5, 681±14 at week 6 and 698±14 at week 7). While the fracture prevention step of the rtFE led to an increase in variation, it still was lower in comparison to all other time points past week 4.

## Discussion

In this study, we demonstrated that the introduction of an adaptive loading approach to individualise load would lead to reduced mechanical environment variance in a femur defect model in mice. Mechanical environments under the assumption of constant loading remain divergent throughout the reparative and remodelling phase of week 3 to week 7. This was more pronounced in the event of substantial bone growth seen in the loaded group, where mechanical stimulation is constantly decreasing under an assumption of constant load for loaded mice. Conversely, a substantial increase in median strain in the control mice is observed with declining bone volume. With the constant loading assumption, the lowest variation in median strain is at week 7, where all bones have sufficiently bridged and fragile structures of the callus are being or have been modelled into new cortical bone or have been removed in the remodelling processes. This indicates how changes in geometry from loading interventions cause large changes in the mechanical environment, which by the very nature of the fracture-healing environment are present, with or without loading. Contrasting the constant loading assumption, when rtFE adaptation is applied, even with fracture prevention screening, we can clearly see that we are able to keep the median strains similar, and reduce group variance, for all mice. The rtFE adaptation responds with changes in bone volume, with control mice requiring a decrease in mechanical load throughout the study, i.e. as excess bone tissue is removed via remodelling, a similar mechanical load would produce a greater median strain within the tissue, hence the load needs to be reduced to compensate for these changes. Contrastingly, the loaded mice display an increasing level of loading required to develop a sufficient strain within the tissue, this is due to loading causing greater bone formation and hence more structural regions to support a particular load. From the displayed data, it is quite clear that if one wants a consistent or targetable degree of mechanical stimulation, one needs to counter the changes in geometry during the repair and remodelling phases, and hence adaptation of loading is required to minimise strain variance between mice at the tissue level. On the other hand, it is clear that using a constant load will lead to varying mechanical environments during the course of fracture healing, delivering drastically different mechanical stimulation to each mouse and its bone tissue. Real time FE adaptation allows the identification of a set of loading parameters (Fig 6), which would achieve a more consistent mechanical environment both longitudinally, throughout the study, as well as cross sectionally within any group, or across groups.

Even though we selected a target distribution with a median strain of 700 microstrain, this in practice can be any choice of distribution centred on any particular median value. However, it would obviously make sense to select a distribution that one hypothesises would give a particularly effect, such as bone growth. In the event of pharmaceutical intervention, one that could aim to create an environment as close to perfect quiescence as possible, minimising the confounding of the pharmaceutical intervention by a longitudinally or cross sectionally varying mechanical environment. Additionally, rtFE has a primary advantage over approaches based on strain gauge measurements on the surface of the bone^35-37^. RtFE allows surfaces, which are inaccessible to strain gauges, yet active in formation and resorption events in the callus region, to be included in stain targeting.

While the fracture prevention approach is conservative, it plays a critical role in preventing load adaptation reaching dangerously high strains in the potential fragile environment within the reparative and remodelling phase. Often, once the time point has been reached where the experimental design calls for loading to begin, there are several animals with partially or incompletely bridged fractures. In figure 4b&c, we can see the difference from a delicate truss-like structure to a more solidified cortical structure within the same bone via large changes in median strain. While the majority of mice for this model heal quickly and well, other models can include larger defects that require more time to bridge^16^, leaving the start date of loading in doubt. Even in the presented data, where the fracture gaps are relatively small, several mice displayed low bone volume at week 3 and hence had less structure to absorb load. This led to small, fragile structures within the callus being over loaded. This can be clearly seen in figure 3a&b, where a very small relative amount of voxels exceed the 1% strain margin, yet in figure 3c, one can observe that many of these highly strained voxels lie next to one another in the thin structural parts of the callus. This increases the risk of refracture and hence poses a risk to the animal. From this visualisation, it becomes clear why the general strain distribution can be misleading in terms of identifying the appropriate load. Without proper visualisation and screening for the upper limit of strains in the tissue, there is a high chance of excessive loading of fragile, small structures within the callus, impairing healing, or at worst threating the welfare of the animal. While the above mentioned fracture risk approach is conservative and will not provide the same accuracy of a validated failure assessment of the bone tissue, it provides a metric for justification of reducing loading forces in the event that animal welfare may be compromised. Hence, a conservative estimate is appropriate given the experimental situation.

Limitations of the above work are largely related to simulation accuracy and correct description of boundary conditions. Micro-FE has been validated as an approach to simulate strains within normal bone as well as callus^38-41^. However, it has proved challenging to validate micro-FE at a tissue level and well specified boundary conditions are essential for accurate results ^41^. Since the core intent of the rtFE adaptation will be implementation in conjunction with a loading machine in an experimental setting, boundary conditions merely need to match such an actuators behaviour. Regardless, this would be a requirement for accurate modelling of the mechanical environment in post-processing and mechano-regulation analysis. This implies that the best use of the concept of adaptive loading is in conjunction with a well-designed and accurate mechanical loading device in defect healing experiments. For this dataset, we assumed only uniaxial loads. However, the boundary conditions could be expanded to any set of uniaxial, shear, bending or torsional boundary conditions with the same principle and analysis applied. Additionally, it is important to note that this approach is not limited to mice. Targeted mechanical stimulus would have the potential to allow induction of specific strain distributions within any bone, fracture or mechanically responsive tissue whether it be in mice, rats, sheep or humans.

Here we analysed mice with the same fixation systems, same sex, operated on by the same surgeon in the same environment, ideally leading to a best-case scenario for external factors that could drive healing differences, yet variances remained. Even though it is well established that tissue level strains are the main driver behind remodelling and healing^2,16^ the vast majority of mechanical interventions do not aim to target particular strains within the tissue directly. However, it is possible that an individualised adaptive loading approach, derived from longitudinal *in vivo* imaging, targeting a specific strain distribution could lead to improved results, reduced confounds and deliver improved outcomes in studies and approaches that have historically produced mixed results. Finally, with the growth in interest and application of personalised medicine the individualised approaches in mice could possibly be translated to humans. In this case, rtFE adaptation would be applicable for mechanical intervention for fractures in patients.

Mechanical environments differ greatly within defect models, even within a group of mice of identical strain, fixation and surgeon. We have shown the need to reduce the inter-animal variance in tissue scale mechanical strains loaded models of bone healing. We propose to do this via optimising each mechanical environment to achieve as similar tissue level strains as possible between individuals within each group. Mechanical environment optimisation provides several benefits within loading models: a) it allows targeting of a strain distribution, providing a specific median strain within the bone tissue, b) it reduces in-group mechanical environment variance, both longitudinally and cross-sectionally, and c) it reduces the probability of refracture in the callus region of the animal. We propose incorporating mechanical environment optimisation into the experimental pipeline; a process, which we have named real time Finite Element (rtFE) adaptive loading. This approach provides a means of reducing mechanical environment variation throughout the post-bridging phrases. Real Time FE can be executed during the conventional imaging-loading pipeline with minimal additional time under anaesthetic for each animal. We believe that when coupled with accurate identification of mechanical dosages required to optimise defect-healing outcome, rtFE adaptive loading can be used to specify and apply these required dosages to homogenise the mechanical environment and reduce variance in both impaired and non-impaired healing cases.

## Materials and Methods

### Study Design

All animal procedures were approved by the authorities (licence number: 36/2014; Kantonales Veterinäramt Zürich, Zurich, Switzerland). We confirm that all methods were carried out and reported in accordance with relevant guidelines and regulations (Swiss Animal Welfare Act and Ordinance (TSchG, TSchV) and ARRIVE guidelines). For the purpose of this study, we used 30 sets of micro-CT images (30 animals, 5 time points per set) taken during a defect healing study in an externally fixated mouse osteotomy model^42^. Each mouse was scanned weekly over a period of 49 days after surgery. 10 of the mice received mechanical loading starting at week 3 post surgery, of which details can be found in^42^. We applied micro-FE analysis to simulate the mechanical environment under constant boundary conditions for all mice, and analysed the longitudinal changes in the mechanical environment of bones. We then applied an adaptive loading approach, targeting the loading to an idealised reference strain distribution, and then analysed and compared the longitudinal changes of the mechanical environment between the constant load dataset and the adaptive load dataset.

### Imaging and pre-processing

All images were acquired by a vivaCT 40 (Scanco Medical, Brüttisellen, Switzerland) with the following scanner settings: 55 kVp, 145 µA, 350 ms integration time, 500 projections per 180°, 21 mm field of view (FOV), aluminium filter. Images were of 10.5 µm resolution. All images were assessed to ensure they were free of artefacts and were of sufficient quality. All original images were cropped to a dimension 300×300×210 voxels and were of the same femoral region between the internal screws of the external fixator for each mouse. The longitudinal images were then registered to the week 0 time-point, to ensure boundary conditions were consistently applied. For post registration, each image was cropped to 180 voxels length via the removal of 15 slices on the top and bottom of the volume. This was done to remove empty slices caused by rotation and translation during image registration. This volume was then Gaussian filtered (σ=1.2, support=1) and we applied the multi-density approach proposed by Tourolle et al ^16^. All grayscale values were binned and then converted from density (mg HA/cm^3^) to Young’s moduli (GPa), on a per voxel basis, from 395 mg HA/cm^3^ to 720 mg HA/cm^3^ in steps of 25 mg HA/cm^3^, corresponding to 4.045 GPa to 12.170 GPa respectively with steps of 0.626 GPa. Regions of soft tissue were set to a Young’s modulus of 0.003 GPa ^15^ and the marrow cavity of the femur was capped with a plate of 20 GPa, preventing edge effects due to the soft tissue found lying on the top slice of the finite element mesh.

### Finite element analysis and scaling of the mechanical environment

From this mesh, the mechanical environment was determined using a linear micro-finite element (micro-FE) solver, Parasol ^43^. A compressive displacement of 1% was applied to the top slice in the axial direction while the bottom slice was fixed. The Swiss National Supercomputing Centre (CSCS) was used to solve each finite element simulation, requiring roughly 2 min per image. Effective strain^44^ results of the simulation were taken as the mechanical environment of the bone.

Two sets of mechanical environments were created from the sets of images (in total 60 mechanical environments per time points (2 per mouse), one at a load of *F*_*applied*_ = 10 N as per Tourolle et al.^16^ and then one at an adaptive load, which was determined by minimising (via a Nedler-Mead optimiser ^45^) the Kolmogorov-Smirnov test result to a reference distribution, determined to be representative of a well-healed mouse with a median strain of 700 microstrain. To align the force on the bone to the constant or calculated values across all mice, the results of the simulations were appropriately scaled based on the assumed loading parameters using the following ratio:

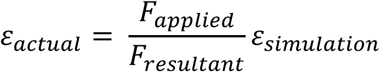

Where *ε*_*simulation*_ is the effective strain result of the simulation (based on the 1% displacement), *F*_*resultant*_is the sum of reaction forces of all the nodes of the uppermost surface, *F*_*applied*_ is the selected force (i.e. a force provided by a mechanical stimulation machine) and *ε*_*actual*_ is the strains under the applied force. All analysis was performed on strains in the bone tissue only, ignoring both the soft tissue and the marrow cap. This was done by masking out regions of soft tissue and marrow caps and then performing all relevant analysis on the remaining voxels.

### Statistics

For each mechanical environment, the median effective strain value was calculated. For all groups at each time point, means and standard deviations were calculated. All histogram were calculated for the range of 0 to 15’000 microstrain with 250 bins over the range. All statistics were performed using Scipy 1.0^45^.

## Acknowledgements

The authors would like to acknowledge support from the European Union (ERC Advanced MechAGE, ERC-2016-ADG-741883). E. Wehrle received funding from the ETH Postdoctoral Fellowship Program (MSCA-COFUND, FEL-25_15-1).

## Author Contributions Statement

The computational study was designed by G.R.P and R.M.. Experimental data was provided from previous *in vivo* studies by G.R.P, E.W. and G.I.K. Data analysis was performed by G.R.P. Additional analysis code was provided by D.C.B. The manuscript was written by G.R.P and reviewed by all authors.

## Data Availability

All necessary data generated or analysed during the present study are included in this published article and its Supplementary Information files (preprint available on BioRxiv). Additional information related to this paper may be requested from the authors.

